# Canopy parkour: movement ecology of post-hatch dispersal in a gliding nymphal stick insect (*Extatosoma tiaratum*)

**DOI:** 10.1101/2020.01.12.903724

**Authors:** Yu Zeng, Sofia W. Chang, Janelle Y. Williams, Lynn Y-Nhi Nguyen, Jia Tang, Grisanu Naing, Robert Dudley

## Abstract

For flightless arboreal arthropods, moving from the understory into tree canopies is cognitively and energetically challenging because vegetational structures present complex three-dimensional landscapes with substantial gaps. Predation risk and wind-induced perturbations in the canopy may further impede the movement process. In the Australian stick insect *Extatosoma tiaratum*, first-instar nymphs hatch on the forest floor and disperse toward tree canopies in the daytime. Here, we address such vertical movements and associated sensory cues in *E. tiaratum* nymphs. Newly hatched nymphs ascend with high endurance, travelling >100 m within 60 minutes. Navigation toward open canopies is underpinned by negative gravitaxis, positive phototaxis, and visual responses to vertically oriented contrast patterns. Nymphal *E. tiaratum* also use directed jumping to cross air gaps, and respond to tactile stimulation and potential threat with a self-dropping reflex, resulting aerial descent. Post-hatch dispersal in *E. tiaratum* thus consists of visually mediated displacement both on vegetational structures and in the air; within the latter context, gliding is then an effective mechanism enabling recovery after predator- and perturbation-induced descent. These results further support the importance of a diurnal niche, in addition to the arboreal spatial niche, in the evolution of gliding in wingless arboreal invertebrates.

**Summary statement:** To effectively disperse into canopies, ground-hatched stick insects use gravity and visual cues to navigate during midday, jump to cross air gaps and respond to threat or perturbation with self-dropping.

## Introduction

Flightless nymphal and adult arthropods compose the majority of arboreal invertebrate fauna (Basset, 2001; Basset et al., 2012); these taxa locomote on vegetational structures (e.g., tree trunks, branches, lianas and leaves) for varied behaviors including locating resources, dispersal, and homing after dislocation (Cunha and Vieira, 2002; Basset et al., 2003; Yoshida and Hijii, 2005). Nevertheless, legged movements by small invertebrates can be challenged by the three-dimensional landscape and dynamic environment of the canopy space. Irregular habitat formed by different vegetational structures (e.g., foliage, branches, and stems) preclude straight displacement, and may also obstruct directed aerial movement within canopy gaps (e.g., forest canopy height ranges globally from 8 – 40 m; Simard et al., 2011). Complexity and dynamics of the physical environment can also impede navigational processes and locomotor performance. Diverse vegetational structures form heterogeneous visual environments under sunlight, with sunflecks and shadows projected dynamically onto various surfaces. The visual environment thus changes with time and with the observer’s perspective, potentially compromising the use of long-range visual cues. Endurance can also be influenced by environmental temperature (e.g., Full and Tullis, 1990), and the daily cycle of temperature in forests (e.g., ~10°C day-night difference in Amazonian rainforest; Shuttleworth, 1985) may constrain the timing and extent of activity. Exposure to predators and parasitoids, many of which are specialized for either searching for or ambushing targets on vegetation (e.g., insects, spiders, and birds; Southwood et al., 1982; Russell-Smith and Stork, 1994; Ford et al., 1986; Basset, 2001) may also threaten invertebrates’ movement through canopy space. In addition to biological threats, environmental perturbations (e.g., rain and wind gusts; see McCay, 2003) can interfere with directed movements. Overall, arboreal transport by small invertebrates can be sensorily and energetically challenging, and such demand may be particularly strong in vertical displacement given the concomitant variation of microhabitats and ecological communities (see Erwin, 1988; Shuttleworth, 1985).

Despite such spatiotemporal variation in the forest environment, both gravity and sunlight can serve as useful long-range cues in canopy space. Terrestrial arthropods generally perceive gravity with gravireceptors and integrate it with other signals (e.g., vision) for navigation (Büschges et al., 2001; Brockmann and Robinson, 2007; Robie et al., 2010). Also, spatial light gradients may be used to infer sun direction in the presence of foliar cover (Bjorkman and Ludlow, 1972; Endler, 1993; Wolken, 1995), with the strongest signal and lowest temporal variation occurring in midday (i.e., a peak plateau of solar altitude with < 5° variation between 11am and 1pm). In addition, vegetational structures under sunlight likely provide short-range visual cues for spatial discrimination. For example, insects identify contrast edges in landing response (Kern et al., 1997; Zeng et al., 2015). The use of gravitational and visual cues is potentially more general than chemical cues, which generally depend on a strong source (e.g., odor from flowers) or permanence (e.g., pheromone trails used by ants; Jackson et al., 2007).

Arboreal invertebrates exhibit varied locomotor behaviors within canopy space. Jumping is a common and energetically efficient strategy for crossing air gaps (see Graham and Socha, 2019). Accidental loss of foothold (e.g., via perturbation by wind or predators) and defensive self-dropping (e.g., startle response: Haemig, 1997; Humphreys and Ruxton, 2019; Dudley and Yanoviak, 2011), along with failed landings after jumping, can transition to directed aerial descent (hereafter termed ‘gliding’), which reduces height loss (see Dudley and Yanoviak, 2011; Yanoviak et al., 2011; Zeng et al., 2016). The diurnal niche for canopy taxa, which permits visually-mediated aerial maneuvers and landing, may thus be essential for the evolution of gliding (Yanoviak et al., 2011; Zeng et al., 2015). Studying the ecology of movement in canopy space, and also initiation mechanisms for aerial descent, can thus help to address the utility of gliding within specific ecological contexts.

The newly hatched nymphs of the Australian stick insect, *Extatosoma tiaratum* Macleay 1826, disperse by ascending from the forest floor, where eggs are deposited, into the canopy (**Fig. 1A,B**). In contrast to the typically nocturnal lifestyle of phasmids, *E. tiaratum* nymphs hatch during midday (i.e., 11 a.m. to 3 p.m.) in the rainy season (November to January; Carlberg, 1981, 1983, 1984; Brock and Hasenpusch, 2009; Brock, 2001), and immediately start to rapidly move. The nymphs also possess ant-mimicking coloration and exhibit comparable behaviors (e.g., body shaking during crawling; **Fig. 1**; **Movie S1**), while climbing and exploring their surroundings (Carlberg, 1981, 1983; Rentz, 1996). These traits collectively form a disperser’s syndrome (Ronce and Clobert, 2012) that potentially facilitates ascent and the search for suitable post-hatch microhabitats. The hatchlings can also glide to reduce height loss if falling (see Zeng *et al*., 2015). Such post-hatch dispersal behavior is exhibited only during the first 3-5 days after hatching, following which the nymphs become nocturnally active and lose their ant-mimicking morphology (**Fig. 1B**) (Brock, 2001; Brock, 1999).

**Figure 1.**
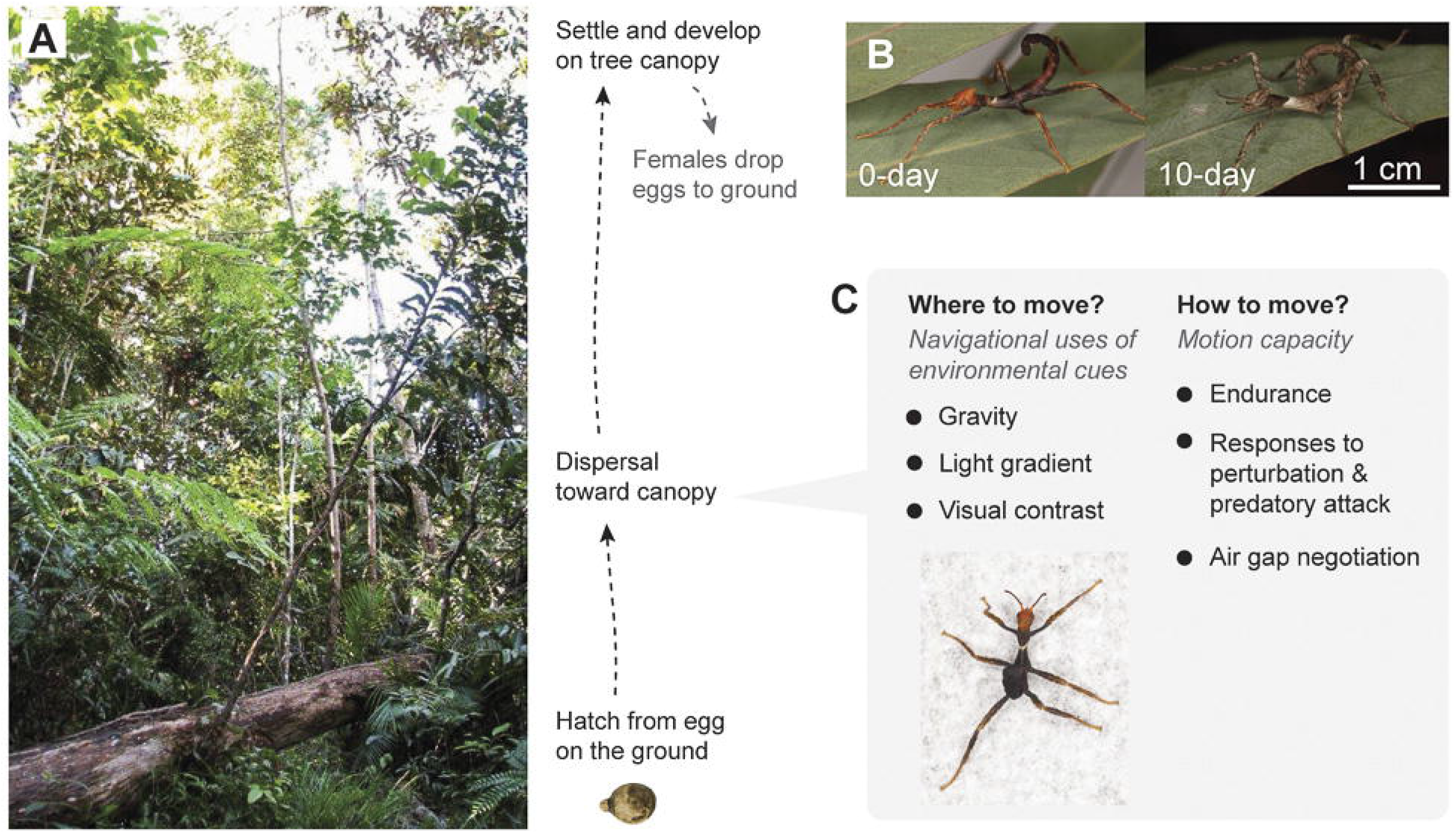
Post-hatch dispersal in *E. tiaratum*. (A) (Left) Natural habitat of *E. tiaratum* illustrating the spatial complexity and vertically varying light environment. Photo was taken in lowland mesophyll forest, Polly Creek, Garradunga, Queensland, Australia (courtesy of Jack W. Hasenpusch). (Right) Schematic summary of *E. tiaratum* life cycle, showing the transient spatial niche of newly hatched nymphs which ascend from forest floor to tree canopies. (B) Loss of ant-mimicking coloration during the first 3 – 5 days after hatching. (C) The two main categories of questions in this study, structured following a ‘movement ecology paradigm’ (see Nathan et al., 2008).

Here, we evaluate movement ecology (Nathan et al., 2008) and canopy dispersal in newly hatched *E. tiaratum* under controlled laboratory conditions, focusing on their navigational and locomotor mechanisms for traveling through the canopy. We hypothesize that: (1) dispersing nymphs use gravity, light gradients, and visual contrast to navigate toward vegetational structures, which are then used for ascent; (2) newly hatched nymphs are capable of considerable vertical ascent; (3) when dispersing, newly hatched nymphs can also initiate aerial descent both volitionally and in response to tactile perturbation. Displacement by individual dispersing insects within vegetational structures across air gaps derives from both locomotor capacity and sensory responses (Fig. 1C), and can explain both movement ecology of small wingless invertebrates in canopy space, and the ecological utility of gliding within the canopy.

## 2. MATERIALS AND METHODS

### 2.1 Experimental insect husbandry

Eggs of *E. tiaratum* were incubated on vermiculite substratum at ~70% humidity and 25–27 °C air temperature. Newly hatched nymphs were collected every 12 hours and were maintained in clear plastic cups (354 ml) with caps. Nymphs were provided with fresh leaves of Himalayan Blackberry *(Rubus armeniacus)* since the day of hatch, and were lightly sprayed with water every 2–3 days. Cups were kept in an environmentally controlled room with a 12h:12h light:dark cycle). All experiments were conducted at air temperatures of 25–27°C.

### 2.2 Negative gravitaxis

A vertically oriented T-maze within a uniformly lit chamber was used. A T-shaped system of wooden rods (diameter, 9 mm) was placed in the center of the experimental chamber (40×40×40 cm), which was covered with white felt and surrounded by natural-spectrum bulbs (R30FF100VLX, Verilux Inc., VT, USA) with voltage control (**Fig. 2A**). Luminance at the center of the chamber was measured with a lightmeter oriented horizontally (Model #401025, Extech, MA, USA; spectral range: 400–740 nm). In each trial, the experimenter first released the insect through the entrance window and then observed the insect’s directional choice through an observation window. A minimum of 3 minutes resting time was given between trials both within and among individuals. Experiments were conducted for three age groups (0–1, 6–7, and 12–14 day-old) under four environmental luminances (0 lx, 10 lx, 600 lx, and 1.5 × 10^4^ lx, adjusted using the voltage controller). Each combination of age and luminance was tested with 10–26 individuals, using 4–6 trials per individual. A repeated G-test (McDonald, 2015) was used to test frequencies of occurrence of negative gravitaxis (f_ascent_) relative to a null frequency of 50%. A Poisson GLMM was used to test whether counts of ascending responses were significantly correlated with environmental luminance and age, using individuals as random factors.

**Figure 2.**
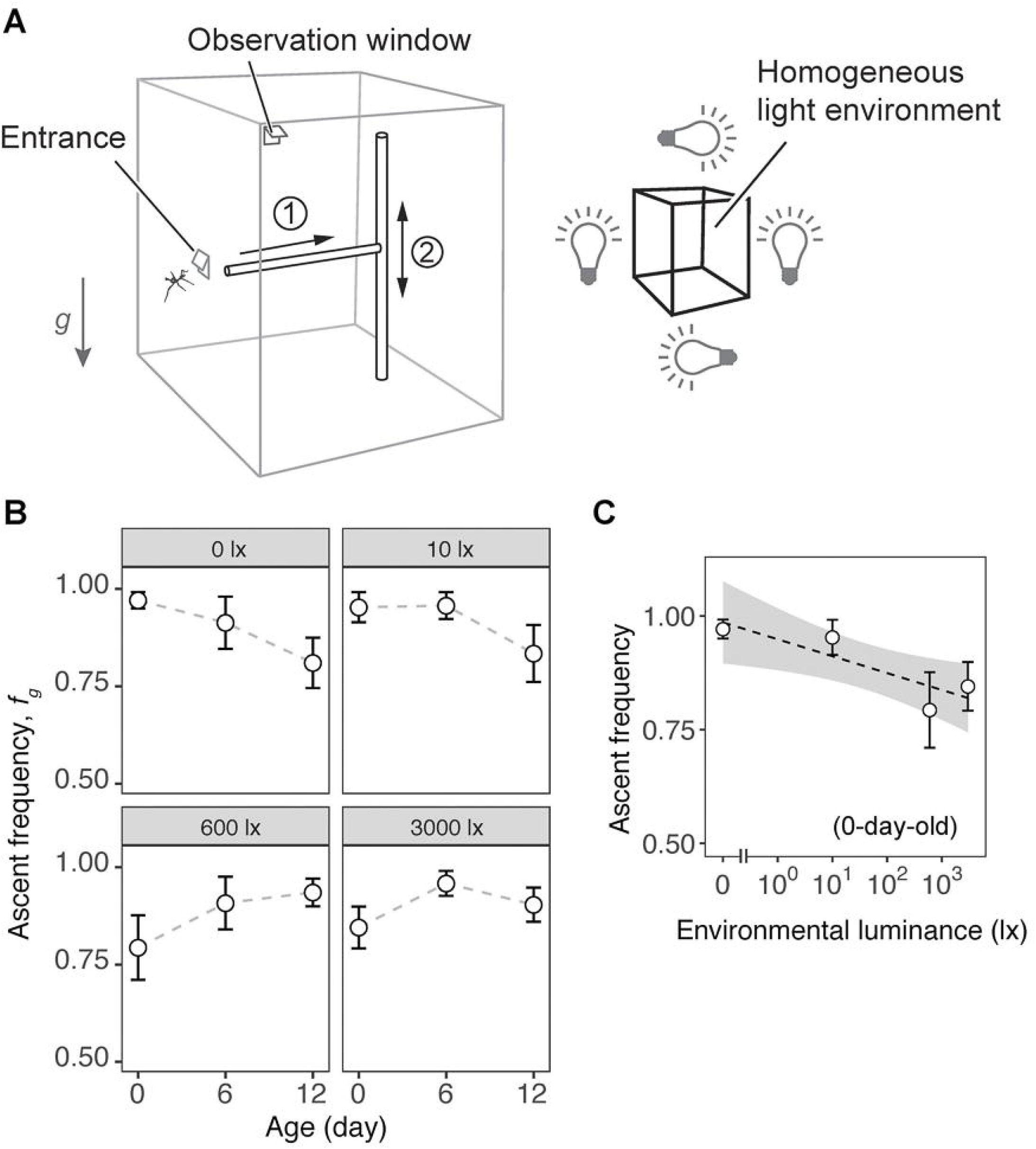
Nymphal *E. tiaratum* exhibited negative gravitaxis under various environmental luminances. (A) Schematic demonstration of the experimental setup. A vertically oriented T-maze made of two rods installed within a cubic chamber covered with white felt, surrounded by voltage-controlled light bulbs for homogeneous luminance (Right). An entrance (2×2 cm) on the chamber wall was opened for the horizontal portion of the T-maze, and an observational window (2×2 cm) was opened at the chamber top. Experimenters first released the insect to the terminal of the horizontal rod (Step 1), and then observed the directional choice of the insect (Step 2). (B) Ascent frequency (*f_g_*) versus age under various environmental luminances, showing a general negative gravitaxis throughout the first instar. Ontogenetic decline of ascent frequency was found in total darkness (0 lx). Values represent means ± s.e.m. All frequencies are significantly different from the null predictions (*P* < 0.001, repeated G-test). See Methods for sample sizes. (C) In newly hatched (0-day-old) nymphs, ascent frequency was inversely correlated with the environmental luminance (Poisson regression coefficient = −0.0013 ± 0.0002, *P* < 0.0001, Poisson GLMM).

### 2.3 Phototaxis

A T-maze was used to test phototactic response of insects. The T-maze consisted of two tunnel sections (section 1: diameter of 2 cm, length 20 cm; section 2: diameter of 4.5 cm, length 10 cm); the interior of both sections was covered with black fabric to reduce light reflection (**Fig. 3A**). A natural-spectrum bulb (R30FF100VLX, Verilux Inc., VT, USA) with a voltage controller was placed ~5 cm away from each exit of the T-maze, with a paper screen (5×5 cm) positioned vertically 3 cm distant from each exit. Luminance was measured with a light meter (401025, Extech, MA, USA; spectral range: 400–740 nm) at two exits. Three luminance contrasts (0 lx vs. 3 lx, 100 lx vs. 500 lx, and 10000 lx vs. 20000 lx) were tested, which range covers the luminance variation between forest understories and canopies (e.g., ~600 lx to 2.4×10^4^ lx; Bjorkman and Ludlow, 1972; Pearcy, 1983; Lee, 1987). Each luminance contrast was tested for insects of three ages (0–1 day-old, 5-day-old, and 10-day-old), each represented by 5–20 individuals (with 5–6 trials per individual). During each trial, the experimenter first released the insect into the entrance, immediately covered the entrance with a lid and then recorded the insect’s directional choice within the light gradient. Control experiments were conducted with one 0–1 day-old group in complete darkness, for which an infrared video camera (SONY HDR-HC7) was used to observe the insects’ directional choice. A repeated G-test was used to analyze significance of directional bias relative to the null frequencies (i.e., 1:1). Poisson generalized linear mixed models (GLMM) were used to test the correlation between counts of phototaxic responses and environmental luminance, using individuals as random factors.

**Figure 3.**
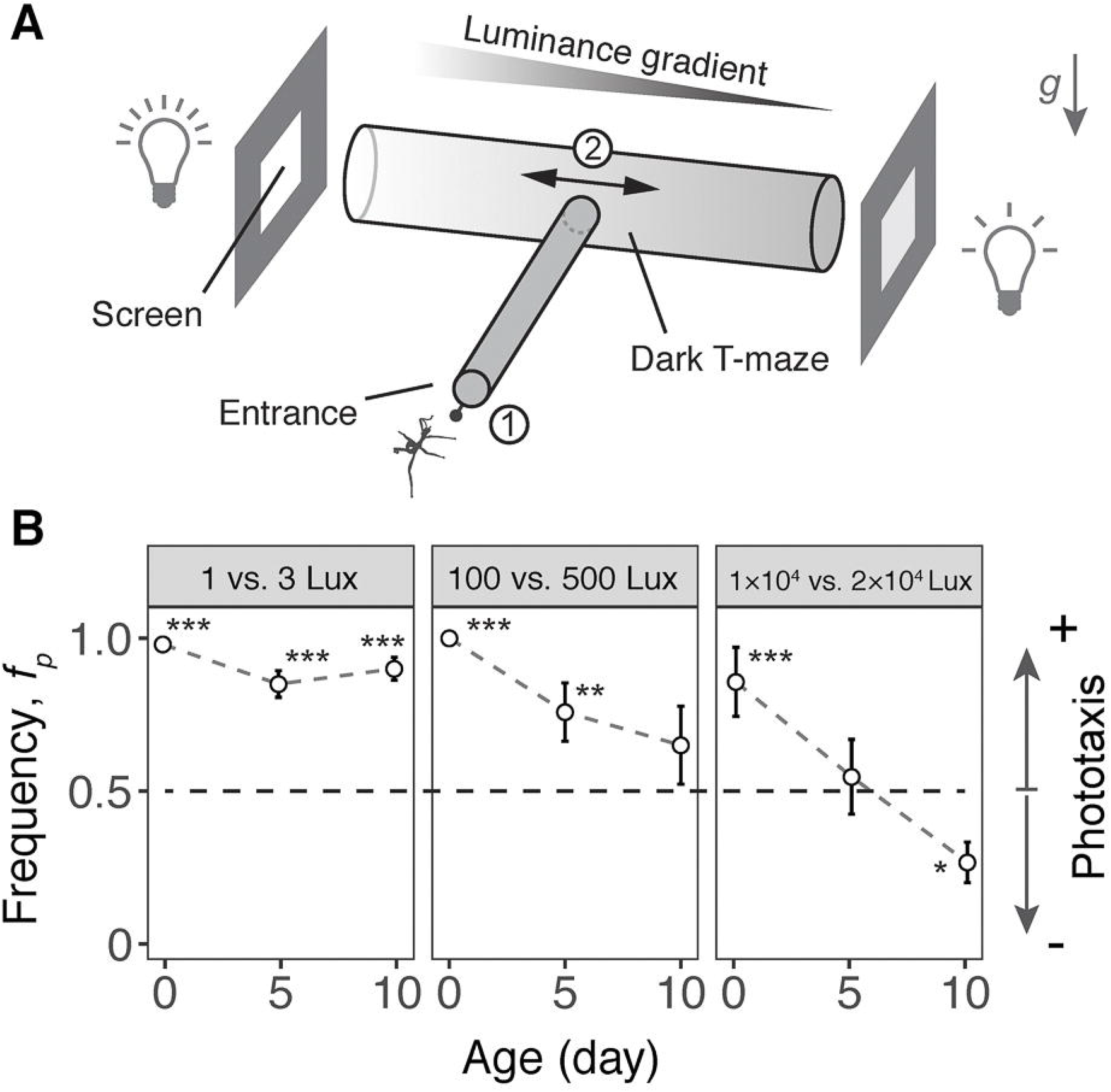
Nymphal *E. tiaratum* exhibited positive phototaxis. (A) Schematic depiction of the experimental setup in a dark room, consisting of a T-maze made of two dark tunnels with a luminance gradient at the intersection. Experimenters first released the insect at the entrance (Step 1), and then recorded its directional choice within the luminance gradient (Step 2). (B) Ontogenetic variation of the phototaxis response under different luminance gradients. Values represent frequencies of phototaxic movements (*f_p_*; means ± s.e.m.). Asterisk symbols denote significance level of results based on repeated G-tests: *, *P* < 0.05; **, *P* < 0.01; ***, *P* < 0.001. In older nymphs, the frequency of phototaxic response was inversely correlated with average luminance (5-day-old, *P* < 0.05; 10-day-old, *P* < 0.01; Poisson GLMM). See Methods for sample sizes.

### 2.4 Tactic responses to visual contrasts

Directional choices were examined using a cylindrical arena (height of 35 cm, diameter 26 cm) decorated internally with vertically oriented contrast patterns. The patterns were made with felt sheets using shades of black, gray, and white (average reflectance over 300–550 nm: black, 3%; gray, 38%; white, 47%), as measured with a spectrophotometer (USB2000, Ocean Optics, Dunedin, FL, USA; see Zeng *et al*., 2015). A natural spectrum light bulb (R30FF75VLX, Verilux Inc., VT, USA) was placed on top of the arena and provide ~600 lx luminance at the arena floor (as measured with a light meter, Extech 401025, Waltham, MA, USA) (**Fig. 4A**).

**Figure 4.**
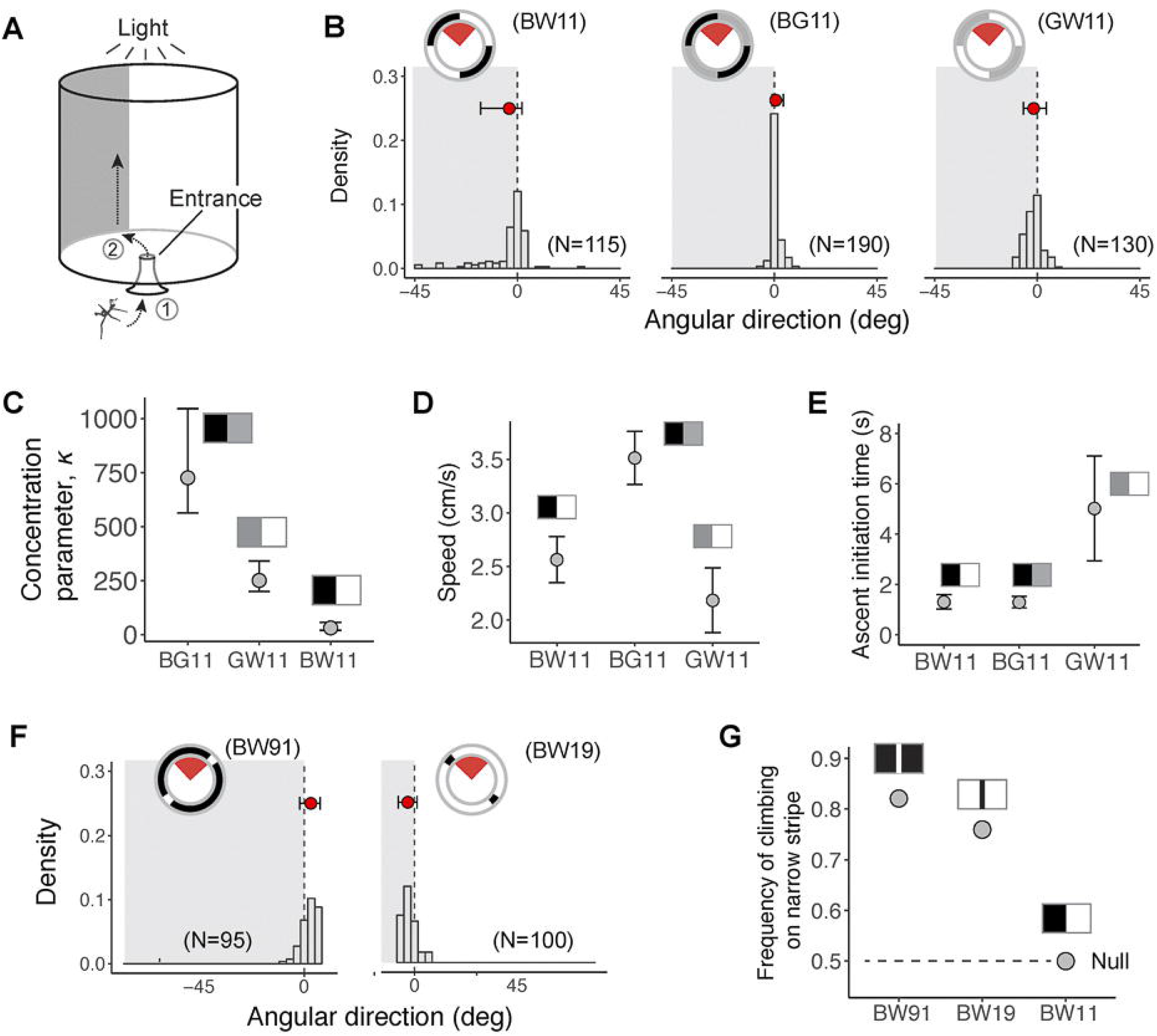
Newly hatched nymphal *E. tiaratum* are attracted by vertically oriented visual contrasts. (A) Schematic illustration of visual arena, the inner wall of which was lined with different contrast patterns (see Methods). Experimenters first released the insect to the arena entrance (Step 1), and then observed its behavior until reaching the arena wall (Step 2). (B) – (C) Summary of directional choices for visual contrasts formed between surfaces of equal size. Subplots in (B) summarize directional preferences with respect to the contrast edge, with histograms illustrating the distributions of movement direction. Insets are schematic diagrams of the visual arena from top view, with the red sectors showing plotted angular ranges. Red dots denote mean angular position, with error bars showing 10th and 90th percentiles. Movement direction exhibited the greatest concentration with a black-gray contrast, as shown by a comparison of the concentration parameter *κ* (means±95% CI) in (C). (D) A comparison of mean speed of movement toward the arena wall with different contrast patterns. The insects moved with highest fastest speed when exposed to the black-gray contrast. Values are significantly different among trials (repeated-measures ANOVA, *F*_2,27_ = 5.09, *P* < 0.05). (E) Comparison of ascent initiation time (the period between reaching the arena wall and initiating ascent) with different contrast patterns. Insects exhibited a significant delay when exposed to gray-white contrast; values are significantly different between trials (*F*_2,27_ = 3.43, *P* < 0.05, repeated-measures ANOVA). For (D) and (E), values represent means±s.e.m. (F)-(G) Summary of directional choices for contrasts formed between surfaces of different sizes. Insects exhibited preference for the narrower surface regardless of its brightness, as shown by significantly greater frequencies relative to the null (G). For (D) and (F), the number of experimental insects and trials were shown in parentheses. See also **Movie S2** and **Supplementary Table S1-S2**; see Methods for sample sizes.

Five contrast patterns were used to test directional preferences to contrast strength and stripe size. The patterns formed by black and white surfaces were coded as BW11, BW91 and BW19, whereby the embedded numbers represent the relative proportions of black (B) and white (W) surfaces. In BW91 and BW19, the narrow stripes feature a spatial frequency of ~3.18 cyc/rad (as viewed from the arena’s center). Similarly, two patterns formed by gray surfaces paired with black or white surfaces in equal proportions were coded as BG11 and GW11. The effect of contrast strength to the insects’ tactic response was tested using BW11, BG11, and GW11, and the effect of stripe size was tested using BW11, BW19 and BW91. Each contrast pattern was tested using 19–38 individuals, and with 5 trials per individual.

In each experimental trial, the insect was released through an entrance (diameter 2 cm) at the center of the arena floor, and was observed from above. Three temporal landmarks were recorded: (1) entrance into the arena, (2) moment of arrival at the arena’s wall, and (3) initiation of ascending. The insect’s directional preference was represented by the angular distance between the point where the insect first reached the arena wall and the nearest contrast edge. Circular directionality of the insects’ directional choice was analyzed using ‘CircStats’ package, which calculates means and confidence intervals of direction *θ* and concentration *κ* based on von Mises Maximum Likelihood Estimates (Lund and Agostinelli, 2007). Repeated G-tests were used to test directional preference for the darker and the lighter surfaces in each configuration against null proportions, which were derived based on random association with corresponding areal proportions of different surface patterns. Durations between temporal landmarks, and also the average speed of movement on the arena floor, were compared using repeated-measures ANOVA.

### 2.5 Ontogenetic decline in ascent endurance

We recorded ascending movements of insects moving on a vertically oriented treadmill (width of 2 cm, height 28 cm), which was connected to a speed controller and placed in a dark room (**Fig. 5A**). A light source was placed above the treadmill to phototactically motivate the insects. A reference disc with visual marks was attached to the top shaft of the treadmill to indirectly indicate treadmill speed. The insect and treadmill were filmed in lateral view using a digital video camera (25 fps; HDR-XR160, Sony Corporation, Japan). For each trial, the experimenter first released the insect onto the track and then maintained the insects’ position within the live video feed by manually adjusting treadmill speed. Nymphs of six ages were used (2, 4, 48, 96, 120, and 240 hour-old; see **Supplementary Table S3**); each individual was filmed for 60 min. Motion tracking software (ProAnalyst, Xcitex, MA, USA) was used to track instantaneous position of the insect and the orientation of the reference disc. Custom-written MatLab scripts were then used to calculate average ascending speed as a function of time.

**Figure 5.**
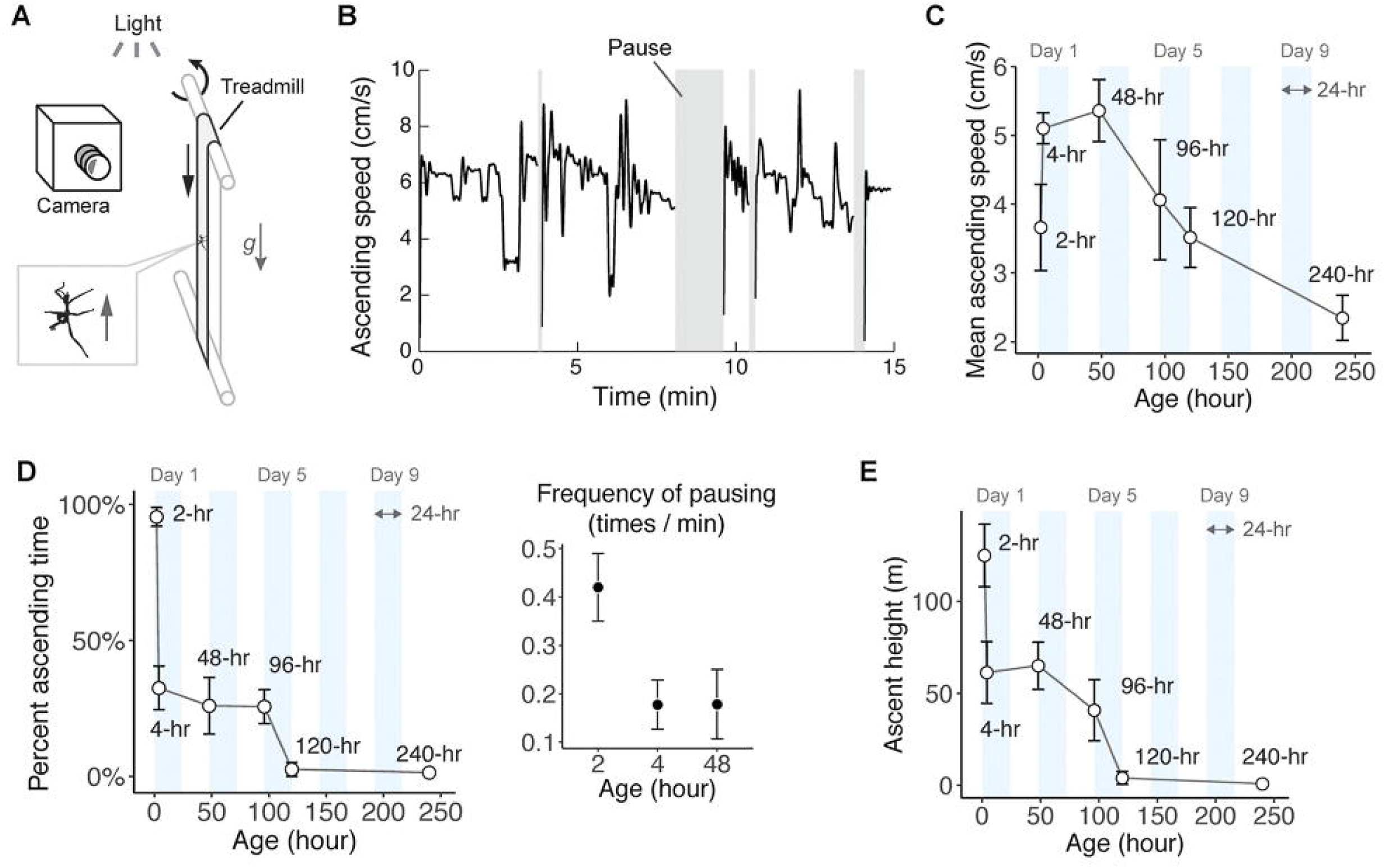
Ontogeny of ascending behavior in nymphal *E. tiaratum*. (A) Schematic of the experimental setup, illustrating a vertically oriented treadmill with a light source on the top. The ascending insect was maintained in camera view via manual speed control by the experimenter. (B) A representative speed profile, showing active ascending movements with intermittent pauses. (C) Ontogenetic variation in ascent speed. Mean speeds are significantly different between age groups (One-way ANOVA, *F*_5,20_ = 18.4, *P*<0.001). (D) The temporal percentage of active ascent relative to the total time, showing a sharp reduction after 2-hr. Inset shows a significantly greater frequency of pausing in newly hatched (2-hr) relative to older nymphs. (E) Comparison of total ascent height within 60 min between different age groups, showing significant inter-group differences (One-way ANOVA, *F*_5,18_ = 6.14, *P*<0.01,). For (B)–(E), values represent means±s.d.; see also **Supplementary Table S3**.

### 2.6 Self-dropping behaviors

Simulated predatory attacks and induction of accidental loss of foothold were applied to experimental insects ascending on a vertically oriented cardboard sheet (width of 15 cm, height, 35 cm) placed beneath a light source. Experimental insects were first released near the bottom of the sheet, and were allowed to ascend under volitional phototaxis and negative gravitaxis. To simulate predatory attacks, a compressed cylinder of paper towel (diameter ~5 mm) was held against the insect’s tibial and tarsal segments for any given leg, pinning it against the substrate for ~20 ms without causing injury (**Fig. 6A**; **Movie S3**). The experiment was conducted for nymphs of three age groups (0-day-old, 6-day-old, and 12-day-old), using 10–15 individuals per group. Each individual was tested in five trials; in each trial, each leg pair received two simulated attacks, with a waiting period of 10 sec between consecutive attacks. To induce accidental losses of foothold, a Teflon-coated surface (~2 cm wide) was introduced to the cardboard sheet (**Fig. 6C**). The insects’ response to contact with this slippery surface was recorded for nymphs of three age groups (0-day-old, 5-day-old and 10-day-old), using 7 individuals per group.

**Figure 6.**
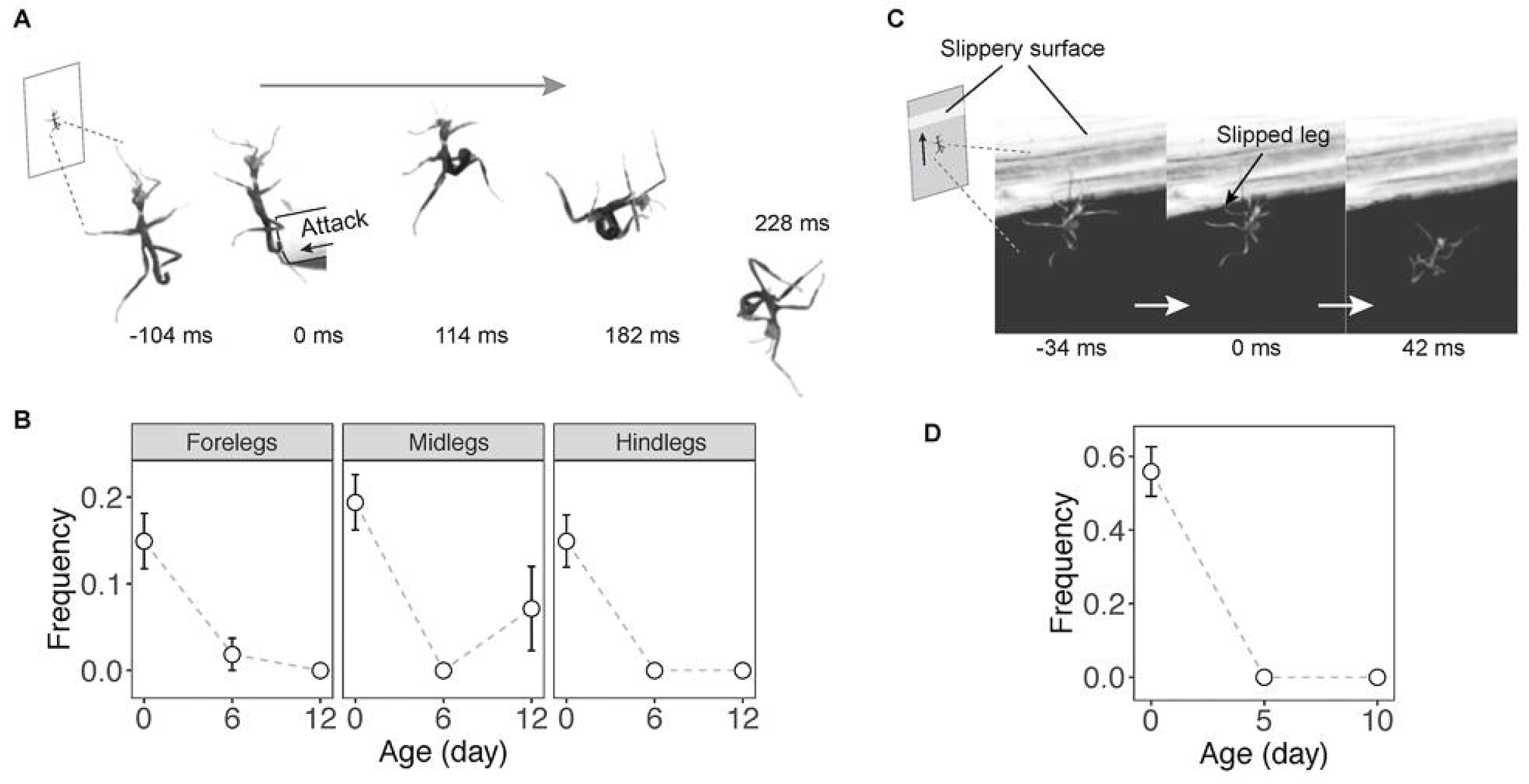
Newly hatched *E. tiaratum* initiate self-drop under tactile stimulations. (A) High-speed video sequence of initiation of self-dropping in response to simulated predatory attack. (B) Newly hatched nymphs showed a significantly higher frequency of self-dropping after simulated attacks. Values represent means±s.e.m. (0-day-old, N = 14 individuals and N = 124 trials per leg pair; 6-day-old, N = 22 insects and N = 208 trials per leg pair; 12-day-old, N = 10 insects and N = 100 per leg pair); see also **Movie S3**. (C) A sample sequence of self-dropping in response to sudden loss of foothold when contacting a slippery surface. The inset depicts the experimental setup. (D) Newly hatched nymphs showed a significant higher frequency of self-dropping after sudden loss of foothold. Values represent means±s.e.m. (0-day-old, N = 7 insects and N = 59 trials; 5-day-old, N = 7 insects and N = 50 trials; 10- day-old, N = 7 insects and N = 50 trials); see also **Movie S4**.

### 2.7 Jumping behavior for crossing air gaps

We evaluated jumping behavior in nymphs by introducing them to an elevated platform (2 cm × 5 cm), which was mounted atop a vertically oriented rod and ~50 cm above the experimental chamber’s floor and > 40 cm away from its walls. The floor and walls were covered with white felt, the environmental luminance was ~600 lx. Two landing targets were used: (1) A horizontally oriented rod (diameter of 5 mm) wrapped with black felt and positioned 20 cm below the platform, and with a white background ~30 cm below the platform (**Fig. 6A**); (2) A vertically oriented rod, also wrapped with black felt, and positioned ~8 cm away from the platform (**Fig. 6A**). For each of these two configurations, jumping behavior was tested for nymphs of three age groups: 0-day-old, 6-day-old, and 12-day-old (horizontal target, 6–9 individuals per age group and 5 trials per individual; vertical target, 5–9 individuals per age group and 2–4 trials per individual). In each trial, the experimenter released the insect onto the vertical rod and then observed its response for ~4 min after it ascended to the platform. If the insect stopped moving, the experimenter gently tapped the vertical rod to stimulate the insect upward. A minimum of 2 min resting period was provided between trials. Control experiments were conducted in five 0-day-old individuals following the same protocols, but without any visual landing targets.

## 3 Results

### 3.1 Negative gravitaxis

After being released into a vertically oriented T-maze within a visually homogeneous environment (**Fig. 2A**), nymphs moved to the intersection and ascended along the vertical rod under various environmental luminances (0 lx – 3000 lx), exhibiting frequencies of gravitaxic response (*f_g_*) significantly different from the null (**Fig. 2B**). Gravity alone can thus be used by nymphal *E. tiaratum* as a directional cue. Furthermore, ascent frequency was inversely correlated with the environmental luminance in newly hatched (0-day-old) nymphs, with the greatest frequency (*f_g_* ~1) occurring in total darkness (**Fig. 2C**).

### 3.2 Positive phototaxis

After being released into a T-maze of horizontal dark tunnels, nymphs moved into a controlled luminance gradient. Three luminance contrasts were used, with the mean value ranging from ~2 lx to 1.5×10^4^ lx (**Fig. 3A**). Newly hatched nymphs (i.e., 0-day-old) exhibited strong phototaxic displacement under all luminance conditions, with the frequency of phototaxic response (*f_p_*) > 0.8 under all light conditions. Overall, *f_p_* declined with both age and increasing mean luminance (**Fig. 3B**). For example, 10-day-old nymphs tended to move toward the darker direction when mean luminance exceeded 1.5 × 10^4^ lx. In control experiments with no luminance gradient, insects showed only non-significant directional bias (repeated G-test: *G* = 0.722, d.f. = 1, *P* = 0.396), supporting the hypothesis that nymphal *E. tiaratum* use luminance gradients alone for directional reference.

### 3.3 Tactic movement toward visual contrasts

Experimental insects were released into a visual arena surrounded by vertically oriented contrast lines (**Fig. 4A**; **Movie S1**). When contrast patterns were formed by equally sized surfaces (black-white, black-gray, and gray-white), insects rapidly moved toward contrast lines (**Fig. 4B**), showing the strongest directional preference to a black-gray contrast (**Fig. 4C**; **Supplementary Table S1**). When exposed to black-gray contrasts, they also exhibited the fastest movement speeds (~3.5 cm/s) on the arena floor, and initiated climbing immediately after reaching the wall (< 2 s) (**Fig. 4D,E**). Black-white and gray-white contrasts were less attractive; the insects showed the slowest movements when exposed to the gray-white contrast. Furthermore, when exposed to contrast patterns formed between surfaces of different sizes, insects showed a general preference for the narrower surface independent of its brightness. For such contrast patterns formed between black and white surfaces, the frequencies of moving toward the narrower surface were significantly greater than the null expectation (**Fig. 4F,G**). The tactic response to vertically oriented contrast lines and preference to narrower surfaces may help in localizing vegetational structures with which to initiate ascent (see Discussion).

### 3.4 Ontogenetic decline of ascent endurance

We evaluated ascending behavior of nymphal *E. tiaratum* for six age groups ranging from 2-hour-old to 240-hour-old individuals. Experimental insects were placed on a vertically oriented and speed-controlled treadmill, and were filmed for 60 min (**Fig. 5A**). Ascents on the treadmill were then digitized to generate speed profiles, consisting of intermittently ascending and pausing intervals (**Fig. 5B**). Ascent speed (range 2.3–5.3 cm/s) generally tended to decline with increasing age. The 4-hour-old and 48-hour-old nymphs exhibited the highest speed, peaking within the first three days after hatching (**Fig. 5C**). A low ascent speed in 2-hour-old nymphs derived from frequent pauses (**Fig. 5D**). The 2-hour-old nymphs exhibited the greatest ascent capacity, including the greatest time spent ascending (> 90% active time) and the greatest total ascent height (100+ m); older (> 5-day-old) nymphs were active for < 1 min and otherwise remained immobile throughout the trial (**Fig. 5D,E**).

### 3.5 Predator- and perturbation-induced self-dropping

Experimental insects were tested with two tactile perturbations, namely simulated predatory attacks and accidental losses of foothold (see Methods). For newly hatched nymphs, simulated attacks led to self-dropping (**Fig. 6A,B**). By contrast, older nymphs generally oriented away from the direction of attack. High-speed filming revealed that self-dropping was initiated by voluntary withdrawal of the tarsus from the substrate within ~50 ms following termination of simulated attack, followed by mid-air tucking (i.e., flexion of tibia-femur joint and elevation of femur) of all leg pairs (N = 30 trials; **Fig. 6A**; **Movie S3**). Slippery surfaces were used to induce loss of footholds during ascent, such that experimental insects were subjected to an unexpected sudden imbalance and loss of foothold (**Fig. 6C**). Newly hatched nymphs (0- day-old) then exhibited self-dropping behavior at a frequency of 55.9 ± 6.7% (mean ± s.e.m.; **Fig. 6D**). High-speed videos revealed that, following loss of foothold by one leg, other leg pairs showed similar flexion and tucking as described above, leading to complete removal of tarsi from the substrate and self-dropping (**Movie S4**). Self-dropping after touching the slippery surface was not observed in older nymphs.

### 3.5 Jumping for crossing air gaps

To test if nymphal *E. tiaratum* jump to cross air gaps, we placed individual nymphs at the base of an elevated platform with a distant visual target (either a horizontal rod or a vertical rod) which contrasted with a white background (**Fig. 7A**). After ascending to the platform, the insects typically oriented toward the visual target, explored the space, and tried to reach out using forelegs. Having failed to cross the air gap using the forelegs, the youngest insects (0-day-old) would then jump toward the visual target, as observed in 19 out of 24 tested individuals (**Fig. 7B**; **Movie S5**). When exposed to the horizontal target, insects spent 1.8±0.2 min (mean ± s.e.m., N = 28 trials from 9 individuals) before jumping and successfully landing on the target in 17 out of 28 trials (success rate ~60%); for trials with successful landing, they traveled a mean horizontal distance of 8.2±1.6 cm with a 20 cm descent (and thus with a ratio of horizontal to vertical distance of ~0.41). When exposed to the vertical target, the insects spent 2.8±0.3 min (N = 21 trials from 9 individuals) before jumping, and successfully landed in 13 out of 23 trials (success rate ~56%); for trials with successful landings, they descended 14.4±6.1 cm with an 8 cm horizontal translation (and a horizontal to vertical distance ratio of ~0.55. No jumping behavior was elicited in control experiments with no visual target (N = 5 individuals). In contrast to the 0-day-old insects, older insects (i.e., 6-day-old and 12-day-old) generally showed no jumping behavior, and either climbed down the vertical rod of the platform or rested in a still posture on the platform after trying to reach out with forelegs.

**Figure 7.**
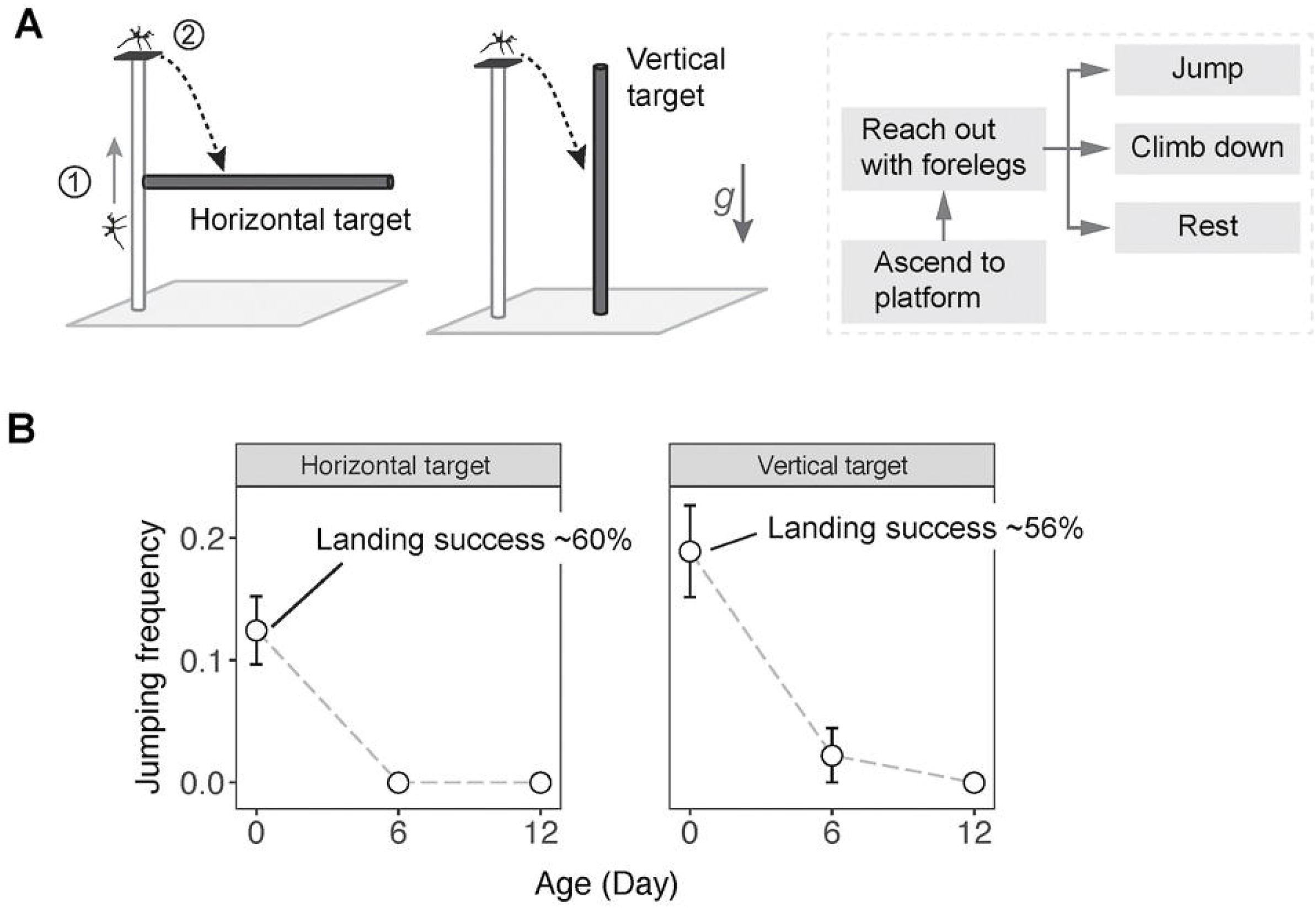
Newly hatched *E. tiaratum* jump to cross air gaps. (A) In each experimental trial, the insect first ascends to the jumping platform through a vertical wooden rod (Step 1), and its subsequent response to a distant land target (dark rod with a white background) was then recorded (Step 2). Inset shows the general behavioral sequence. (B) Variation in jumping frequency versus insect age, showing significant ontogenetic reduction in jumping behavior. Values represents means±s.e.m. See Results for statistical summary; see also **Movie S5**.

## 4. Discussion

### Canopy dispersal and the evolution of gliding in *E. tiaratum*

The tactic uses of gravity and environmental light gradients by newly hatched *E. tiaratum* support our initial hypotheses on the roles of these two cues for legged movement in canopy space, as found in other ecosystems (e.g., other larval insects, and mites; Perkins et al., 2008; Zhang, 1992). Tactic movement toward vertically oriented contrast lines suggests searching behavior for vertical structures, especially stems and tree trunks, for use in ascent. Preference for dark or shaded surfaces may reduce exposure to visual predators during dispersal; a similar preference was shown in gliding nymphal stick insects (Zeng et al., 2015). This preference nonetheless persists under low luminance (e.g., <5 lx; **Fig. 3B**), suggesting high visual acuity under low light. To further understand cognitive abilities underlying dispersal in nymphal *E. tiaratum*, studies of their movements within structurally and visually more complex environments would be informative. The potential use of other directional cues (e.g., polarized light, chromatic signals, chemical cues, etc.) in the canopy space can be tested using binary choice experiments. Recording movement on natural vegetation (e.g., via a Lagrangian approach; Baguette et al., 2014) can provide direct evidence for integrative use of various short-range and long-range cues.

A multimodal locomotor strategy, consisting of both aerial and legged phases, can assist small insects moving in canopy space. We predict that, under natural conditions, dispersing *E. tiaratum* may frequently alternate movements on vegetational structures and in air, thus forming a behavioral loop (**Fig. 8A,B**). An aerial phase may be initiated by missed landings, environmental perturbations, and predatory attacks. After dropping, energy required for subsequent ascent can be indirectly reduced by aerial righting and gliding, rather than falling to the ground. The self-dropping reflex that triggers the withdrawal of all tarsi may present a reflex arc similar to avoidance reflexes at the single-leg level (Kittmann, 1996). Foothold withdrawal may underlie a variety of voluntary self-dropping mechanisms in phasmids (Bedford, 1978) and some other insects (e.g., ants and aphids; see Haemig, 1997; Humphreys and Ruxton, 2019).

**Figure 8.**
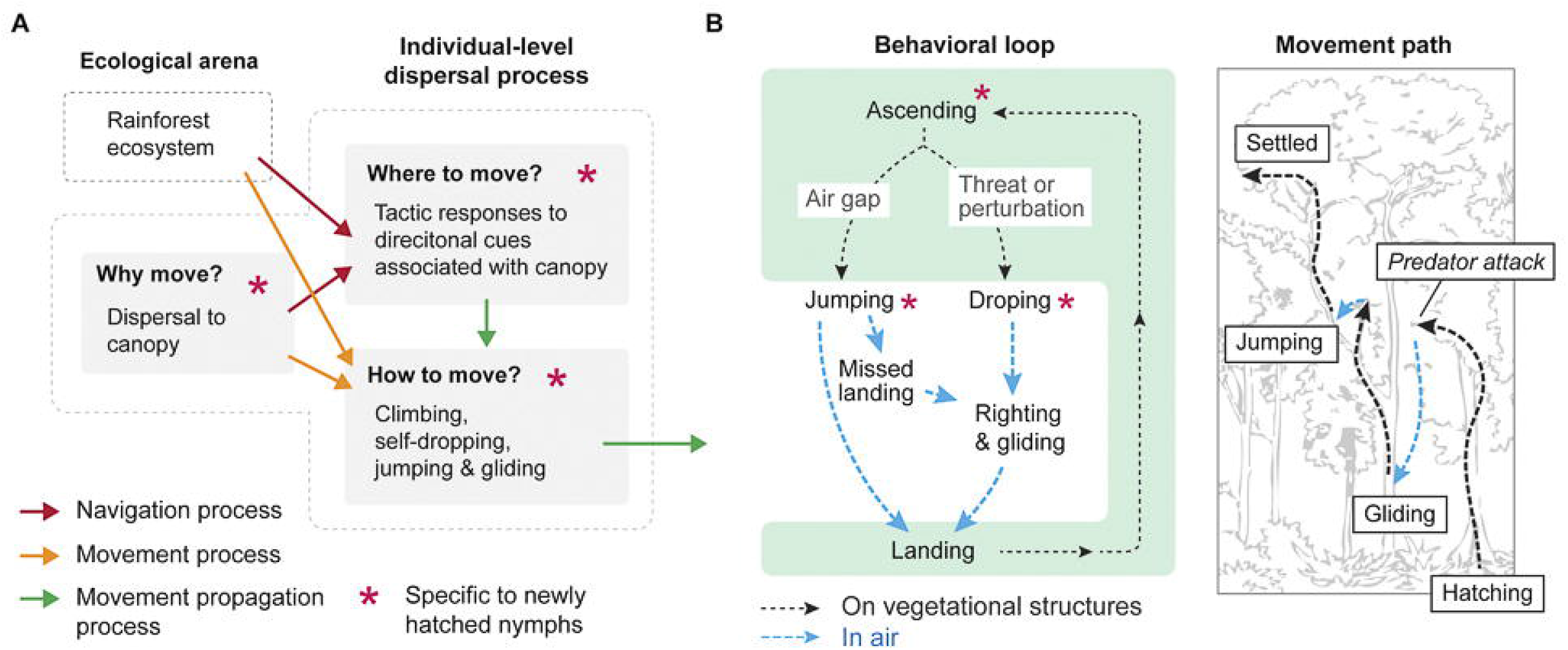
Ecology of canopy dispersal and gliding in newly hatched *E. tiaratum*. (A) Navigation and movement processes of post-hatch dispersal in *E. tiaratum*, whereby alternation between movements on vegetational structures and in air form a behavioral loop (B), as shown by a schematic (right).

The diurnal niche plays a key role in gliding behavior of nymphal *E. tiaratum.* Control of midair maneuvering and landing depends on the strength and continuity of vertically oriented contrast edges (e.g., tree trunks; see Zeng et al., 2015), the qualities of which decreases with and would be less salient at lower light levels. Compound eyes of arthropods generally possess much lower acuity compared to vertebrate eyes (Land, 1997), and will be more dependent in flight on visual contrast associated with tree trunks. Therefore, a diurnal niche may be essential to the evolution of gliding in arthropods, and perhaps less so in vertebrates. This conclusion is indirectly supported by the observation of widespread diurnal gliding in various arthropods (see Dudley and Yanoviak, 2011; Yanoviak et al., 2015), in contrast to crepuscular and nocturnal gliding in various vertebrate gliders (e.g., colugos; Byrnes et al., 2011). Spatiotemporal variation of visual quality of landing targets within vegetational canopies has not yet been examined relative to arthropod gliding, but clearly is relevant to evolution of this behavior. Similarly, few data pertain to the occurrence and behavioral contexts of aerial descent (e.g., as elicited as a defensive strategy or in response to environmental perturbation) in a range of arboreal invertebrates (see Yanoviak et al., 2011; Yanoviak et al., 2015).

### Evolution of diurnal dispersal in nymphal phasmids

Phasmids are generally nocturnal (Bedford, 1978), whereas diurnal hatching in *E. tiaratum* likely evolved under selection for rapid dispersal within a rainforest ecosystem. Ascent of the ground-hatched nymphs is a transient behavior that utilizes environmental cues for navigation and movement. As environmental brightness is strongest around midday (Shuttleworth, 1985), hatching around this time maximizes use of this cue (see **Fig. 9A**). Furthermore, ant mimicry (i.e., myrmecomorphy) by newly hatched *E. tiaratum* may help to deter visual predators during movement on vegetational structures. Similar ant-mimicking phenotypes are described from newly hatched nymphs in several unrelated phasmid species (**Fig. 9B**; see also Hanibeltz et al., 1995). Nymphs of all such taxa hatch on the ground, and are diurnally active. Convergent evolution of this strategy implies a common demand for dispersal efficiency, particularly given the high abundance of nocturnally active predators on vegetation (see Berger and Wirth, 2004). Hatching location, diurnality, and vegetational features underlie thus evolution of diurnal ascent in nymphal phasmids, although a broad phylogenetic survey is now warranted to evaluate possible correlates of other dispersal strategies.

**Figure 9.**
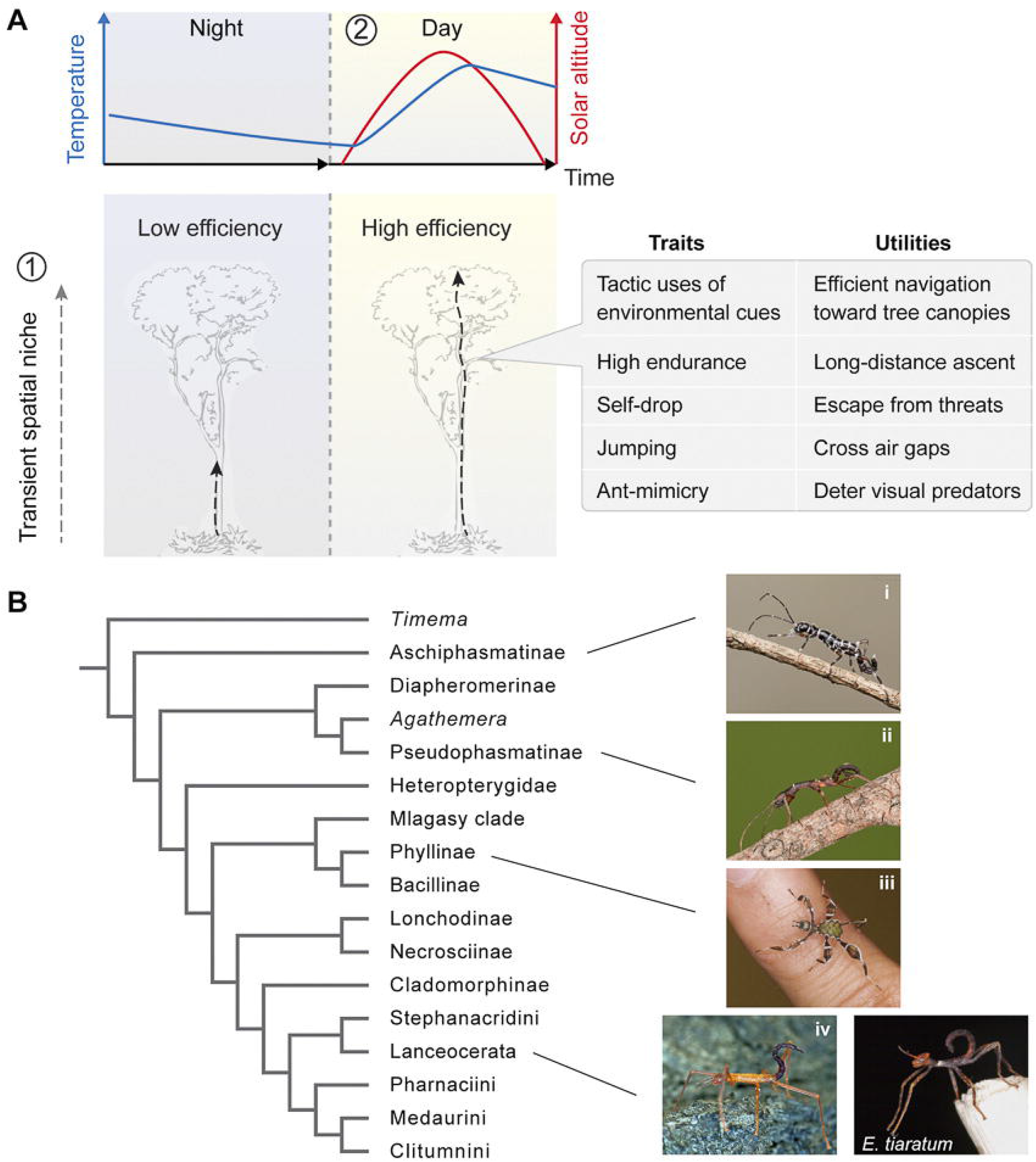
Evolution of diurnal dispersal in newly hatched stick insects. (A) Diurnal dispersal in nymphal *E. tiaratum* is likely an evolutionary consequence of the interplay between (1) an obligatorily transient spatial niche, and (2) a diurnal temporal niche under selection for greater dispersal efficiency. (B) Examples of newly hatched nymphs that are ground-hatched and diurnally active with ant-mimicking coloration and behavior, mapped on a recent phylogeny of main phasmid clades (Simon et al., 2019). (i) *Orthomeria* sp. (Aschiphasmatidae) from Sarawak, Borneo (photo courtesy of Bruno Kneubühler); (ii) *Paraprisopus* sp. (Prisopodidae) from Panama (photo courtesy of Bruno Kneubühler), with phylogenetic position based on that of a related clade (Prisopodini); (iii) *Phyllium philippinicum* (Phylliidae) from Philippines; (iv) *Podacanthus* sp. (Phasmatidae) from Queensland, Australia (photo courtesy of Tony Eales).

## Acknowledgments

We thank Douglas Ewing for inspiration, and David Wake and members of Berkeley Animal Flight Laboratory for comments. We also thank Andrea Lucky for communication about *Leptomyrmex* ants, and Paul Brock and Jack Hasenpusch for advice on *E. tiaratum* natural history. We further acknowledge research facilities provided by the Center for Interdisciplinary Biological Inspiration in Education and Research (CiBER), UC-Berkeley. This work was partially funded by the Undergraduate Research Apprentice Program (URAP) of UC-Berkeley.

## Author contributions

Y.Z. devised the experiments. Y.Z., S.C., J.W., L.N., J.T. and G.N. participated in data collection and data analyses. All authors contributed to writing the manuscript.

## Supplementary Movies

S1 – Newly hatched *E. tiaratum* nymphs dispersing; note ant-mimicking coloration and movements.

S2 – Representative trial of a *E. tiaratum* nymph navigating within the experimental arena with visual contrasts.

S3 – Self-dropping reflex after a simulated attack, shown at 5x reduced speed (original video filmed at 500 fps).

S4 – Self-dropping reflex in an *E. tiaratum* nymph following contact with a Teflon-coated surface.

S5 – Representative trials of jumping behavior used to cross air gaps.

## Symbols and abbreviations

GLMM: generalized linear mixed model
*θ*: circular direction of von Mises distribution
*κ*: concentration parameter of von Mises distribution

## Notes

### Competing Interest Statement

The authors have declared no competing interest.

## References

Baguette, M., Stevens, V. M. and Clobert, J. (2014). The pros and cons of applying the movement ecology paradigm for studying animal dispersal. Mov Ecol 2, 13.

Basset, Y. (2001). Invertebrates in the canopy of tropical rain forests How much do we really know? Plant Ecol. 153, 87–107.

Basset, Y., Cizek, L., Cuénoud, P., Didham, R. K., Guilhaumon, F., Missa, O., Novotny, V., Ødegaard, F., Roslin, T. and Schmidl, J. (2012). Arthropod diversity in a tropical forest. Science 338, 1481–1484.

Basset, Y., Kitching, R., Miller, S. and Novotny, V. (2003). Arthropods of tropical forests: spatiotemporal dynamics and resource use in the canopy, Cambridge University Press.

Bedford, G. O. (1978). Biology and ecology of the Phasmatodea. Annu. Rev. Entomol. 23, 125–149.

Berger, J. R. and Wirth, R. (2004). Predation-mediated mortality of early life stages: a field experiment with nymphs of an herbivorous stick insect *(Metriophasma diocles*). Biotropica 36, 424–428.

Bjorkman, O. and Ludlow, M. M. (1972). Characterization of the light climate on the floor of a Queensland rainforest. Carnegie Inst. Wash. Yearb. 71, 85–94.

Blaesing, B. and Cruse, H. (2004). Stick insect locomotion in a complex environment: climbing over large gaps. J. Exp. Biol. 207, 1273–1286.

Brock, P. (1999). The Amazing World of Stick and Leaf Insects, AES Publishing.

Brock, P. D. (2001). Studies on the Australasian stick-insect genus *Extatosoma* Gray (Phasmida: Phasmatidae: Tropoderinae: Extatosomatini). J. Orthopt. Res. 10, 303–313.

Brock, P. D. and Hasenpusch, J. W. (2009). Complete Field Guide to Stick and Leaf Insects of Australia, CSIRO Publishing.

Brockmann, A. and Robinson, G. E. (2007). Central projections of sensory systems involved in honey bee dance language communication. Brain. Behav. Evol. 70, 125–136.

Büschges, A., Schmidt, J. and Wolf, H. (2001). Sensory processing in invertebrate motor systems. e LS.

Byrnes, G., Libby, T., Lim, N. T. L. and Spence, A. J. (2011). Gliding saves time but not energy in Malayan colugos. J. Exp. Biol. 214, 2690–2696.

Byrnes, G., Lim, N. T. L., Yeong, C. and Spence, A. J. (2011). Sex differences in the locomotor ecology of a gliding mammal, the Malayan colugo *(Galeopterus variegatus*). J. Mammal. 92, 444–451.

Cant, J. G. (1992). Positional behavior and body size of arboreal primates: a theoretical framework for field studies and an illustration of its application. Am J Phys Anthropol 88, 273–283.

Carlberg, U. (1981). Hatching-time of *Extatosoma tiaratum* (Macleay) (Phasmida). Entomol.’s Mon. Mag. 117, 199–200.

Carlberg, U. (1983). A review of the different types of egglaying in the Phasmida in relation to the shape of the eggs and with a discussion on their taxonomic importance (Insecta). Biol. Zent.bl. 102, 587–602.

Carlberg, U. (1984). Hatching Rhythms in *Extatasoma tiamtum* (MacLeay) (Insecta: Phasmida). Zool. Jahrb., Abt. allg. Zool. Physiol. Tiere. 88 (4), 441–446.

Carlberg, U. (1984). Oviposition behavior in the Australian stick insect *Extatosoma tiaratum*. Cell. Mol. Life Sci. 40, 888–889.

Cunha, A. A. and Vieira, M. V. (2002). Support diameter, incline, and vertical movements of four didelphid marsupials in the Atlantic forest of Brazil. J. Zool. 258, 419–426.

Dudley, R. and Yanoviak, S. P. (2011). Animal aloft: the origins of aerial behavior and flight. Integr. Comp. Biol. 51, 926–936.

Endler, J. A. (1993). The color of light in forests and its implications. Ecol. Monogr. 63, 2–27.

Erwin, T. L. (1988). Biodiversity. In Biodiversity (ed. E. O. Wilson), pp. 123–129. National Academies Press.

Evans, G. C. and Coombe, D. E. (1959). Hemisperical and woodland canopy photography and the light climate. J. Ecol. 47, 103–113.

Ford, H. A., Noske, S. and Bridges, L. (1986). Foraging of birds in eucalypt woodland in north-eastern New South Wales. Emu 86, 168–179.

Full, R. J. and Tullis, A. (1990). Capacity for sustained terrestrial locomotion in an insect: energetics, thermal dependence, and kinematics. J. Comp. Physiol. B 160, 573–581.

Graham, M. and Socha, J. J. (2020). Going the distance: The biomechanics of gap-crossing behaviors. J. Exp. Zool. A: Ecol. Genet. Physiol. 333, 60–73.

Haemig, P. D. (1997). Effects of birds on the intensity of ant rain: a terrestrial form of invertebrate drift. Anim. Behav. 54, 89–97.

Hanibeltz, A., Nakamura, Y., Imms, A. and Abdullah, E. (1995). The survival of newly-hatched leaf insects. Phasmid Studies 4, 60.

Humphreys, R. K. and Ruxton, G. D. (2019). Dropping to escape: a review of an under-appreciated antipredator defence. Biol Rev Camb Philos Soc 94, 575–589.

Jackson, D. E., Martin, S. J., Ratnieks, F. L. and Holcombe, M. (2007). Spatial and temporal variation in pheromone composition of ant foraging trails. Behav. Ecol. 18, 444–450.

Kern, R., Egelhaaf, M. and Srinivasan, M. V. (1997). Edge detection by landing honeybees: behavioural analysis and model simulations of the underlying mechanism. Vision Res. 37, 2103–2117.

Kittmann, R., Schmitz, J. and Büschges, A. (1996). Premotor interneurons in generation of adaptive leg reflexes and voluntary movements in stick insects. J. Neurobiol. 31, 512–531.

Land, M. F. (1997). Visual acuity in insects. Annu. Rev. Entomol. 42, 147–177.

Lund, U. and Agostinelli, C. (2007). CircStats: Circular statistics, from “Topics in Circular Statistics” (2001). S-plus original by Lund, U. Rport by Agostinelli, C. Rpackage version 0.2-3

McCay, M. G. (2003). Winds under the rain forest canopy: the aerodynamic environment of gliding tree frogs. Biotropica 35, 94–102.

Nathan, R., Getz, W. M., Revilla, E., Holyoak, M., Kadmon, R., Saltz, D. and Smouse, P. E. (2008). A movement ecology paradigm for unifying organismal movement research. PNAS 105, 19052–19059.

Perkins, L. E., Cribb, B. W., Hanan, J., Glaze, E., Beveridge, C. and Zalucki, M. P. (2008). Where to from here? The mechanisms enabling the movement of first instar caterpillars on whole plants using *Helicoverpa armigera* (Hübner). Arthropod-Plant Inte. 2, 197–207.

Rentz, D. C. (1996). Grasshopper country: the abundant orthopteroid insects of Australia, University of New South Wales Press.

Robie, A. A., Straw, A. D. and Dickinson, M. H. (2010). Object preference by walking fruit flies, *Drosophila melanogaster*, is mediated by vision and graviperception. J. Exp. Biol. 213, 2494–2506.

Ronce, O. and Clobert, J. (2012). Dispersal syndromes. Dispersal Ecol. Evo. 155, 119–138.

Russell-Smith, A. and Stork, N. E. (1994). Abundance and diversity of spiders from the canopy of tropical rainforests with particular reference to Sulawesi, Indonesia. J. Trop. Ecol. 10, 545–558.

Shuttleworth, W. J. (1985). Daily variations of temperature and humidity within and above Amazonian forest. Weather 40, 102–108.

Simard, M., Pinto, N., Fisher, J. B. and Baccini, A. (2011). Mapping forest canopy height globally with spaceborne lidar. J. Geophys. Research-Biogeo. 116.

Simon, S., Letsch, H., Bank, S., Buckley, T., Donath, A., Liu, S., Machida, R., Meusemann, K., Misof, B. and Podsiadlowski, L. (2019). Old World and New World Phasmatodea: Phylogenomics Resolve the Evolutionary History of Stick and Leaf Insects. Front. Ecol. Evol. 7, 345.

Southwood, T. R. E., Mow, V. C. and Kennedy, C. E. J. (1982). The assessment of arboreal insect fauna: comparisons of knockdown sampling and faunal lists. Ecol. Entomol. 7, 331–340.

Wolken, J. J. (1995). Light detectors, photoreceptors, and imaging systems in nature, Oxford University Press, USA.

Yanoviak, S. P., Dudley, R. and Kaspari, M. (2005). Directed aerial descent in canopy ants. Nature 433, 624–626.

Yanoviak, S. P., Munk, Y. and Dudley, R. (2011). Evolution and ecology of directed aerial descent in arboreal ants. Integr. Comp. Biol. 51, 944–956.

Yanoviak, S. P., Munk, Y. and Dudley, R. (2015). Arachnid aloft: directed aerial descent in neotropical canopy spiders. J. R. Soc. Interface 12, 20150534.

Yanoviak, S. P., Silveri, C., Stark, A. Y., Van Stan, J. T. and Levia Jr, D. F. (2017). Surface roughness affects the running speed of tropical canopy ants. Biotropica 49, 92–100.

Yoshida, T. and Hijii, N. (2005). Vertical distribution and seasonal dynamics of arboreal collembolan communities in a Japanese cedar (*Cryptomeria japonica* D. Don) plantation. Pedobiologia 49, 425–434.

Zeng, Y., Lam, K., Chen, Y., Gong, M., Xu, Z. and Dudley, R. (2017). Biomechanics of aerial righting in wingless nymphal stick insects. Interface Focus 7, 20160075.

Zeng, Y., Lin, Y., Abundo, A. and Dudley, R. (2015). Visual ecology of directed aerial descent in first-instar nymphs of the stick insect *Extatosoma tiaratum*. J. Exp. Biol. 218, 2305–2314.

Zhang, Z. (1992). Phototactic and geotactic responses in *Allothrombium pulvinum* larvae (Acari: Trombidiidae). Exp. Appl. Acarol. 15, 41–47.

