## Supplementary Information for "Canopy parkour: movement ecology of post-hatch dispersal in a gliding nymphal stick insect (*Extatosoma tiaratum*)"

### Supplementary material

#### Taxic responses to visual contrast

| | | von Mises distribution | | | repeated G-test with William's correction | | |
|---|---|---|---|---|---|---|---|
| setup | N | $\mu$ (95%CI) | $\kappa$ (95%CI) | $f_D$ (%) | $G_{total}$ | $G_{pool}$ | $G_{heterogeneity}$ |
| BW11 | 100 | -0.06 (6.19, 6.25) | 32.38 (20.4, 57.3) | 57.1 | 35.8 (23 d.f., *) | 2.0 (1 d.f., 0.158) | 33.8 (22 d.f., 0.051) |
| BG11 | 188 | 0.01 (0.01, 0.02) | 727.07 (563.47, 1046.45) | 37.7 | 60.4 (38 d.f., *) | 7.42 (1 d.f., **) | 53.0 (37 d.f., *) |
| GW11 | 127 | -0.03 (6.25, 6.27) | 251.96 (200.78, 340.94) | 75.7 | 72.2 (26 d.f., ***) | 30.6 (1 d.f., ***) | 41.6 (25 d.f., *) |

**Table S1.** Statistical results of movement direction in visual contrast experiments. $f_D$, the frequency of moving towards the darker surface in each setup, excluding those overlapped with contrats edges. G-values are presented with d.f. and $P$-values in parentheses (*, $P$ <0.05; **, $P$ <0.01; ***, $P$ <0.001). See Fig. 4.

| | | | repeated G-test with William's correction | | |
|---|---|---|---|---|---|
| setup | N | $f_{stripe}$ (%) | $G_{total}$ | $G_{pool}$ | $G_{heterogenity}$ |
| BW91 | 95 | 82 | 43.48 (9 d.f., **) | 20.08 (1 d.f., ***) | 23.4 (18 d.f., 0.18) |
| BW19 | 100 | 76 | 86.49 (20 d.f., ***) | 61.04 (1 d.f., ***) | 25.45 (19 d.f., 0.15) |

**Table S2.** Statistical results of movement direction in visual contrast experiments. $f_D$, the frequency of moving towards the darker surface in each setup, excluding those overlapped with contrats edges. G-values are presented with d.f. and $P$-values in parentheses (*, $P$ <0.05; **, $P$ <0.01; ***, $P$ <0.001). See Fig. 4.

#### Ontogeny of ascent activity

| age | N | ascent speed (cm/s) |
|---|---|---|
| 2-HAH | 2 | 3.7±0.9 |
| 4-HAH | 4 | 5.1±0.4 |
| 48-HAH | 4 | 5.4±0.9 |
| 96-HAH | 6 | 4.1±2.1 |
| 120-HAH | 4 | 3.5±0.9 |
| 240-HAH | 4 | 2.3±0.7 |

**Table S3.** Ascending speeds of sampled age groups.
